# An Extended Maximum Likelihood Inference of Geographic Range Evolution by Dispersal, Local Extinction and Cladogenesis

**DOI:** 10.1101/038695

**Authors:** Champak R. Beeravolu, Fabien L. Condamine

## Abstract

The origin and evolution of species’ ranges remains a central focus of historical biogeography and the advent of likelihood methods based on phylogenies has revolutionized the way in which range evolution has been studied. A decade ago, the first elements of what turned out to be a popular inference approach of ancestral ranges based on the processes of Dispersal, local Extinction and Cladogenesis (*DEC*) was proposed. The success of the *DEC* model lies in its use of a flexible statistical framework known as a Continuous Time Markov Chain and since, several conceptual and computational improvements have been proposed using this as a baseline approach. In the spirit of the original version of *DEC*, we introduce *DEC eXtended* (*DECX*) by accounting for rapid expansion and local extinction as possible anagenetic events on the phylogeny but without increasing model complexity (*i.e*. in the number of free parameters). Classical vicariance as a cladogenetic event is also incorporated by making use of temporally flexible constraints on the connectivity between any two given areas in accordance with the movement of landmasses and dispersal opportunity over time. *DECX* is built upon a previous implementation in C/C++ and can analyze phylogenies on the order of several thousand tips in a few minutes. We test our model extensively on Pseudo Observed Datasets and on well-curated and recently published data from various island clades and a worldwide phylogeny of Amphibians (3309 species). We also propose the very first implementation of the *DEC* model that can specifically account for trees with fossil tips (*i.e*. non-ultrametric) using the phylogeny of palpimanoid spiders as a case study. In this paper, we argue in favour of the proposed improvements, which have the advantage of being computationally efficient while toeing the line of increased biological realism.

## Introduction

Historical biogeography attempts to address how the *“earth and life have evolved together, accounting for current geographic patterns in terms of past events”* (Donoghue 2014). In keeping with this wide definition, the main challenge in studying the biogeographic history of a set of species lies in integrating all the factors influencing species ranges which include various geological, climatic, ecological and chance events (Lomolino et al. 2010). Traditionally, research in the this field has been descriptive with the main objective being the reconstruction or estimation of ancestral areas using approaches such as cladistic biogeography (Morrone and Carpenter 1994; Donoghue and Moore 2003). Only recently have inferences based on rigorous statistical frameworks, such as maximum likelihood (hereafter ML) and Bayesian approaches, been used for testing alternative hypotheses of geographical range evolution (Ree and Sanmartín 2009; see the review of Ronquist and Sanmartín 2011). In contrast to former *pattern-based* methods, recently developed parametric, or *process-based*, approaches hypothesize and test for macroevolutionary processes mainly by making use of time-calibrated phylogenies (Ree and Sanmartín 2009). Some of the well known ones include *Dispersal-Vicariance Analysis (DIVA*, Ronquist 1997), *Dispersal-Extinction-Cladogenesis (DEC*, Ree and Smith 2008), and several more recent ones, *GeoSSE* (Goldberg et al. 2011), *SHIBA* (Webb and Ree 2012), *BayArea* (Landis et al. 2013) and *DEC+J* (Matzke 2014). For the sake of clarity, a glossary is provided in **Table 1** and keeps track of the different names, definitions, and objects used in historical biogeography.

**Table 1.** A glossary of the commonly used terms in historical biogeography.

| **Term** | **Synonym (or abbreviation)** | **Definition** |
| --- | --- | --- |
| Adjacency matrix | Connectivity matrix, Areas allowed, Evolving Geological Connectivity | A matrix indicating the presence (1) or absence (0) of a connection between any two areas, and which determines the total number of ranges in the analysis |
| Area | Geographic operational unit | The smallest geographic unit used in a discrete area biogeographic analysis which can vary from an island to a continent |
| BayArea | BayArea (Landis et al. 2013; https://code.google.com/p/bayarea/) | A variant of the DEC model which implements an MCMC exploration of biogeographic histories. It also includes a distance-dependent mode of dispersal (implemented in C++) |
| DEC | Dispersal-Extinction-Cladogenesis (Ree et al. 2005; Ree and Smith 2008; Smith 2009; http://www.reelab.net/lagrange/configurator/index) | The original biogeographic model using a likelihood framework (works under Python and C++) |
| DEC+J | Dispersal-Extinction-Cladogenesis + jump event (Matzke 2014; https://cran.r-project.org/web/packages/BioGeoBEARS/index.html) | A variant of the DEC model which implements a third parameter (j) for modelling the relative effects of founder events at cladogenetic events (works under the R environment) |
| DECX | Dispersal-Extinction-Cladogenesis Extended (this study) | A version of the DEC model in which the classical vicariance and the rapid anagenetic change are included (implemented in C++) |
| Dispersal rate matrix | Dispersal matrix, Rate matrix, Manual dispersal multipliers | A matrix of scaling factors representing the dispersal intensity (or dispersal probability) between two areas (a value between 0 and 1) |
| Distance matrix | Distance-dependent dispersal matrix | A matrix of scaling factors representing the dispersal intensity (or dispersal probability) between two areas as the function of the distance separating them (a value between 0 and 1) |
| DIVA | Dispersal-Vicariance Analysis (Ronquist 1997; http://sourceforge.net/projects/diva/) | A parsimony approach similar to DEC in which anagenetic events are range expansion and extinction, and cladogenetic events are the classical vicariance (works under C++, and implemented in S-DIVA, and RASP) |
| Jump dispersal | Founder-events associated with cladogenetic events | It occurs when a small group of migrants (in this context species), which is not genetically representative of the population from which they came, establish in a new area and ultimately become a new species by allopatry |

**Table 1 (continued)**
|  |  |  |
| --- | --- | --- |
| Lagrange | Likelihood Analysis of Geographic RANGE Evolution<br>(https://code.google.com/p/lagrange/) | The general statistical maximum likelihood frame in which the DEC model has been developed |
| Long-distance dispersal | Transoceanic dispersal | A dispersal described as long to very long distance dispersal often invoked in clades that have colonized remote continents from other continents (e.g. Australia to Madagascar) |
| Q matrix | Transition probability matrix | A matrix of instantaneous transition rates between geographic ranges (or states) |
| Range | Collections of predefined areas, State | One or several areas making an ancestral distribution or current distribution |
| Rapid anagenetic change | Range shifts/Range expansions | In the DECX model, this anagenetic event represents both a partial or complete shift to a new geographic range and to locally go extinct in an ancestral range |
| Rapid cladogenetic change | Jump dispersal | In the DEC+J model, this cladogenetic event represents both a dispersal event to a new area (not present in the ancestral area) plus a local extinction of the daughter lineage |
| Range contraction | Local extinction | In biogeography, this represents an extinction in a given geographic area during the anagenesis |
| Range expansion | Dispersal | In biogeography, this represents a movement towards a new area that is successfully colonized, and thus expands the distribution of the lineage |
| RASP | Reconstruct Ancestral State in Phylogenies (Yu et al. 2015<br>http://mnh.scu.edu.cn/soft/blog/RASP/) | A graphical interface and a platform for inferring ancestral state using S-DIVA (Statistical dispersal-vicariance analysis), Lagrange (DEC), Bayes-Lagrange (S-DEC), BayArea and BBM (Bayesian Binary MCMC) method |
| S-DIVA | Statistical-Dispersal-Vicariance Analysis (Yu et al. 2010;<br>http://mnh.scu.edu.cn/soft/blog/SDIVA/) | A graphical interface and a platform that complements DIVA, implements the methods of Nylander et al. (2008) and Harris and Xiang (2009), and determines statistical support for ancestral range reconstructions |
| SHIBA | Simulated Hisotrical Biogeography Analysis (Webb and Ree 2012;<br>http://phylodiversity.net/shiba/) | A very similar version of the DEC model but in which the approach simulates lineage movement in a discrete spatial and temporal model |
| Stepping-stone dispersal | Inter-island dispersal | A dispersal described as short to medium distance dispersal often invoked in island clades that have colonized an archipelago jumping from island to island (e.g. Galapagos Archipelago) |

**Table 1 (continued)**
|  |  |  |
| --- | --- | --- |
| Time slices | Time interval, time period | A period of time in which an adjacency matrix and a rate matrix are used to represent the dispersal pathways and the intensity of dispersal between two areas according to paleogeography at that time |
| Time-stratified analyses | Constrained analyses (opposite is unconstrained analyses) | A DEC model in which the time-scale of the tree is divided into at least two time slices |

Of all phylogeny based methods, the most successful has undoubtedly been the ML framework *DEC* (Ree et al. 2005; Ree and Smith 2008), which models ancestor-descendant range evolution based on stochastic dispersal and local extinction. The main reason for the success of *DEC* being its versatile statistical framework which allows for it to incorporate various types of spatial and temporal information as constraints, such as from fossils, sea levels, climate, and continental drift (e.g. Clark et al. 2008; Clayton et al. 2009; Mao et al. 2012; Wen et al. 2013; Chacón and Renner 2014; Meseguer et al. 2015). Furthermore, these features have been the basis for the development of several biogeographic models (e.g. *GeoSSE, SHIBA, BayArea* and *DEC+J*) with the aim of answering some of the theoretical and/or computational shortcomings of the *DEC* approach.

For instance, *GeoSSE* reciprocally interlinks lineage diversification and range evolution by introducing area specific speciation rates (Goldberg et al. 2011). While *GeoSSE* is a considerable improvement over the assumptions of *DEC*, the inherent increase in model complexity restricts its use to not more than two areas (see also Goldberg and Igić 2012). In another attempt, the program *SHIBA* implements a forwards-in-time rejection sampling approach which enables it to easily simulate biologically realistic and complex lineage movement, defined by a matrix of physical distances between land units and the effect of their sizes on local extinction rates (Webb and Ree 2012). Although *SHIBA* models dispersal more realistically than *DEC*, inference of ancestral states based on rejection sampling appears to be highly inefficient in such a complex geographic setting. The *BayArea* model, akin to the previous idea of physical distances, extends ancestral range inference to situations which involve a very large number of areas (potentially thousands) by adopting a Markov Chain Monte Carlo (MCMC) exploration of biogeographic histories (Landis et al. 2013). However, this increased capacity comes at the price of a highly restrictive model of geographic speciation (sympatry irrespective of the ancestral range size) and a static landmass configuration throughout the timescale of the phylogeny. More recently, the *DEC+J* model using another parameter *J* adds a new kind of cladogenetic event, associated with instantaneous reproductive isolation arising from founder events (due to rare or long-distance dispersal), than those previously considered by the *DEC* model (Table 1 in Matzke 2014). By adding this third degree of freedom, the *DEC+J* model has been shown to outperform the standard *DEC* model for many island clades (Matzke 2014) but also for more globally distributed groups (e.g. Hosner et al. 2015).

However, an important and yet unaddressed drawback in the original *DEC* model (including the *DEC+J* model) resides in its range inheritance assumptions which excludes *classical vicariance* or allopatry (see Fig. 4a in Ronquist and Sanmartín 2011) as a possible cladogenetic event. The current version of the *DEC* model only allows for situations when speciation is restricted to a very local scale which translates into one of the daughter lineages covering no more than a single area even if its ancestral range was widespread (*i.e*. covering several areas). Ree et al. (2005, p. 2302) have justified this exclusion from the range inheritance scenario by stating that classical vicariance *“invokes a historical area event without considering the spatial and temporal context of that event, and we wish to avoid making this kind of ad hoc hypothesis”*. Hence, without *a priori* knowledge of the connectivity between the areas constituting the ancestral range one cannot attribute biologically relevant ranges to daughter lineages for vicariant events. However, ignoring the spatial and temporal context within a species range, as in the *DEC* model, can also lead to equally irrelevant choices. For instance, let us assume that the true ancestral range at a given node consists of five areas (say A, B, C, D and E) and were connected as beads along a chain (**Fig. 1**). The current assumptions of *DEC* (e.g. alloperipatric speciation) would allow it to consider the two daughter range splits (A-B-C | E) and (**D**), where the “|” represents a discontinuity of the dispersal corridor which makes for an unconvincing scenario on a macroevolutionary scale (**Fig. 1**). Instead, a more plausible scenario would be when the ancestral range (A-B-C-D-E) splits into (A-B-C-D) for one daughter lineage and (E) in the other lineage (under alloperipatric speciation) or into (A-B) for one lineage and (B-C-D) for the other (under classical vicariance). However, these assumptions which are lacking in *DEC* could be essential, especially for ancient clades that might have thrived on wide land masses (e.g. Pangea, Gondwana), which are currently isolated as continents (e.g. Mao et al. 2012).

**Figure 1.**
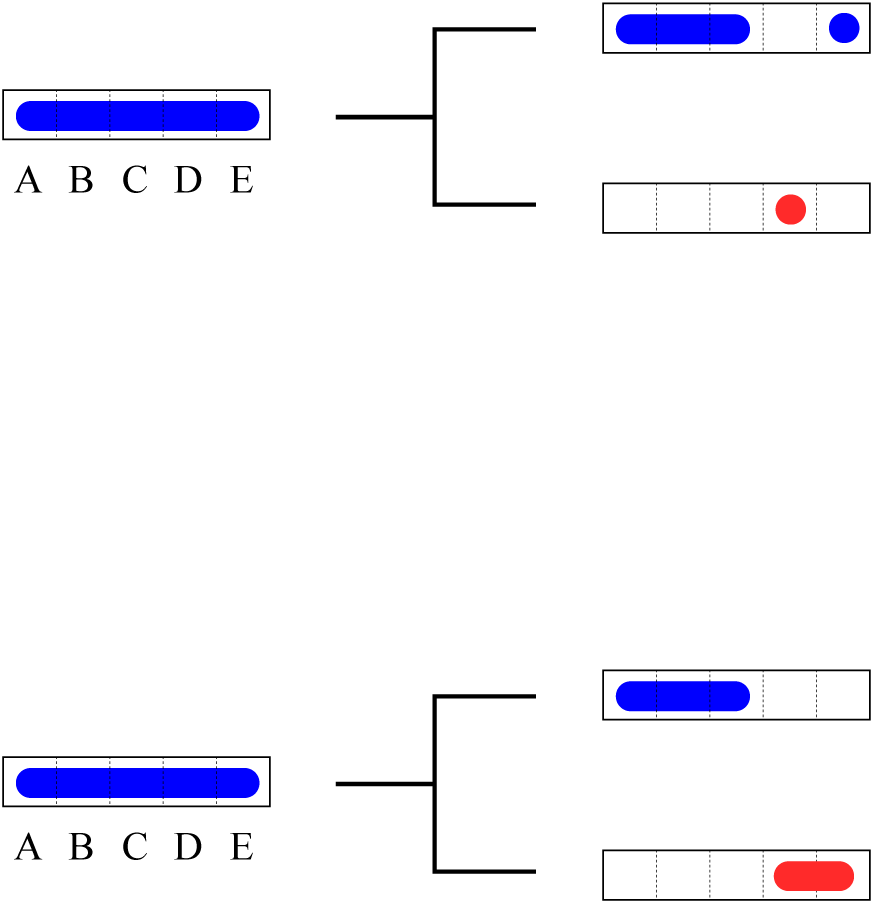
Illustrating the possibility of *the DEC* model to consider spatially incoherent modes of geographic evolution at speciation events on a phylogeny (see also Fig. 4a in Ronquist and Sanmartín 2011). The red and blue colours represent species identities, the bold rectangles represent the entire region of study and the dotted lines within delimit land units or areas which can be grouped to form a species range. *Top*:, inheritance of spatially incoherent daughter ranges at a cladogenetic event; *Bottom*: a classical vicariance scenario.

Another important and missing component of *DEC* lies in its time-stratified model (see **Table 1** for a definition). In its current version, the time-stratified model accounts for temporal changes of dispersal rates among the component areas included in the model, but does not correctly account for tectonic evolution through time (*i.e*. the adjacency matrix is constant over time). Yet, studies have shown that the connectivity/adjacency matrix is paramount in explaining the historical biogeography of a given clade (Buerki et al. 2011; Chacón and Renner 2014). This effect may be even more crucial for ancient clades when the information contained in the form of extant geographic ranges is of less help at deep phylogenetic timescales. We thus extend the original *DEC* framework by allowing for an evolvable range connectivity (evolvable range adjacency) also referred to in this paper as Evolving Geological Connectivity (see **Table 1**). Apart from increased biological and paleogeographic realism, evolving geological connectivity can help significantly reduce the computational burden of inference especially when the biogeographic context contains a large number of areas (Buerki et al. 2011).

Moreover, during anagenesis (geographic evolution along the phylogenetic branches), *DEC* assumes that transitions between geographic states (*i.e*. between ranges) require only a single event, either dispersal towards a previously unoccupied and available area or to go locally extinct in an area within the species range. However, this may seem restrictive given our knowledge on biotic movement in the face of severe climate events (e.g. glaciations; Gavin et al. 2014), or competition with other species/clades (e.g. Alexander et al. 2015). It is biologically plausible that a lineage can partially (or entirely depending on the size of its range) shift its geographic range which would resemble a rapid sequence of events such as dispersal followed by local extinction or *vice versa* (Penz et al. 2015). We refer to this type of geographic evolution as rapid anagenetic change (hereafter *RAC*) which differs from rapid cladogenetic change as modelled by the *J* parameter in the *DEC+J* model.

Despite the increase in the number and complexity of models the paucity of data is of great hindrance, especially for very ancient clades, in the field of historical biogeography. While fossils can come to the rescue and are increasingly being relied upon to infer geographic range evolution, they are still primarily used as *dating tools*. The two major techniques are *node dating* (Ho and Phillips 2009), where fossils are used as calibration points on an ancestral branch and the *tip dating* or *total-evidence* approach (Ronquist et al. 2012) which makes use of a morphological basis to accommodate fossils as extinct tips alongside extant taxa. Both these approaches have also inspired a similar use in historical biogeography and an example of the former (*i.e*. node constraints) can be found in Meseguer et al. (2015). Recently, several others have made use of trees with fossil tips (Mao et al. 2012; Nauheimer et al. 2012; Wood et al. 2013) while relying on the implementations of the *DEC* model as made available by the original authors, either in Python (https://github.com/rhr/lagrange-python) or in C++ (https://github.com/blackrim/lagrange). Both these implementations can lead to an erroneous inference of ancestral ranges when used with non-ultrametric trees. In the following, we provide a remedy to this problem by specifically accounting for trees with fossil tips.

Improvements in this paper are based on an efficient implementation of the *DEC* model (Smith 2009) and address the issues of (*i*) classical vicariance (hereafter *VI*), (*ii*) evolving geological connectivity (hereafter EGC), and (*iii*) rapid anagenetic change (hereafter *RAC*). Our model will hereafter be referred to as the *DEC eXtended* model (or *DECX* in the text); the details of which are presented in the next section. Very importantly, the *DECX* model does not change the total number of free parameters with respect to the original *DEC* model (two parameters). We assess the performance of *DECX* on Pseudo Observed Datasets (hereafter PODs), which are generated for a wide range of lineage diversification scenarios and over an evolving biogeographic context with eight areas. In the following sections, we also apply *DECX* over a number of published datasets on island clades (Matzke 2014) and on a large amphibian phylogeny with over 3300 species (Pyron 2014) for which results from the *DEC* and *DEC+J* models are available. Finally, the phylogeny of palpimanoid spiders (Wood et al. 2013) is used as a case study accounting for trees with fossil tips.

## Methods

### Outline of the DECX model

We provide here but a brief summary of the underlying *DEC* model which has been amply discussed elsewhere (Ree and Smith 2008; Ree and Sanmartín 2009; Ronquist and Sanmartín 2011; Landis et al. 2013). The basic premise of DECX follows closely that of the *DEC* model (Ree and Smith 2008) in that range evolution is modeled on a phylogeny where ancestral ranges (at the interior nodes) are inferred with the help of the geographic ranges of extant species (*i.e*. at the tips). A species range here is an approximation of the underlying continuous geography which is subdivided into several discrete and non-overlapping land units or areas (see Buerki et al. 2011, Fig. 1) based on criteria which can vary from *ecological tolerance* on a local scale to tectonic plate movement on a global level. Note that the sum of these land units makes up the total area of study. Toeing the logic of character evolution models, the geographic states (or ranges) of the extant or ancestral species are the observed or inferred *presence-absence vectors* composed of binary values for these land units respectively.

Mathematically, geographic evolution along the branches of a phylogeny in *DEC* is modeled as a *time-homogeneous* Continuous Time Markov Chain (hereafter CTMC) whose *states* (a fixed number) are geographic ranges consisting of all combinations of the previously defined land units. The choice of a CTMC is warranted by the fact that, “*dispersal and extinction are modeled as Poisson processes with waiting times between events in an area distributed exponentially according to their respective rate parameters*” (Ree et al. 2005, p. 2300). Every Markov model requires the definition of a transition matrix and for a CTMC, the continuous nature of the model demands a matrix of instantaneous rates of change between states (**Q**) (**Table 1**). In the *DEC* model, **Q** is calculated on the basis of a user-defined matrix (**D**) containing the dispersal rates between any two given land units. The latter can either be *a priori* information based on the *aggregate dispersal ecology* of the species or simply some function of the distance matrix for all land units as modelled by *SHIBA* and *BayArea*. The role of the two free parameters *d* and *e* of the *DEC* model is to scale the elements of **Q** depending on whether the change of state is a range expansion or contraction respectively (see Eq. 1 in Ree and Smith 2008). Note that in the DECX model (as in the simplest version of *DEC*), the local extinction rate is assumed to be the same for all land areas. For the sake of computational simplicity, *DEC/DECX* also allow for an upper bound to the number of land units that can constitute a species range which further reduces the total number of states considered for ancestral range inference.

Under the CTMC, the probability of changing from one state to another over a branch of length *t* is given by the matrix of instantaneous probabilities *P(t)* = *e^Qt^*, where **P** is the matrix exponential of **Q**. As pointed out by Landis et al. (2013, p. 791), matrix exponentiation can be computationally onerous when considering a large number of land units. For any given point in the parameter *[d, e]* space, using the **P** matrix and the Felsenstein (1981) pruning algorithm, we are able to calculate the likelihood of a biogeographic history. The last step consists of summing the *conditional probabilities* of the root node being in all of the states. Subsequently, a search of *[d, e]* space using an optimization algorithm yields the ML estimates for *d* and *e* and the associated biogeographic scenario. Ranges at ancestral nodes are further optimized for the cladogenetic process by locally conditioning on the parental and daughter lineages (Ree and Smith, 2008).

For range evolution along the phylogenetic branches (anagenesis), the original *DEC* model considers range expansion/contraction in terms of dispersal from one area to another (parameter *d*) and local extinction within an area (parameter *e*). The *DEC* model makes an assumption that this change in geographic states is limited to the increase/decrease of a species range by a single land unit at every time step. In this paper we improve on this assumption by introducing *RAC* (**Fig. 2**). The biological case for *RAC* arises from possible species responses to severe climate events (e.g. glaciations) which can resemble a shift of its geographic range due to a rapid sequence of events (Gavin et al. 2014, Fig. 1). We also account for other variations of *RAC* which are plausible, such as rapid dispersal or extinction (Penz et al. 2015).

**Figure 2.**
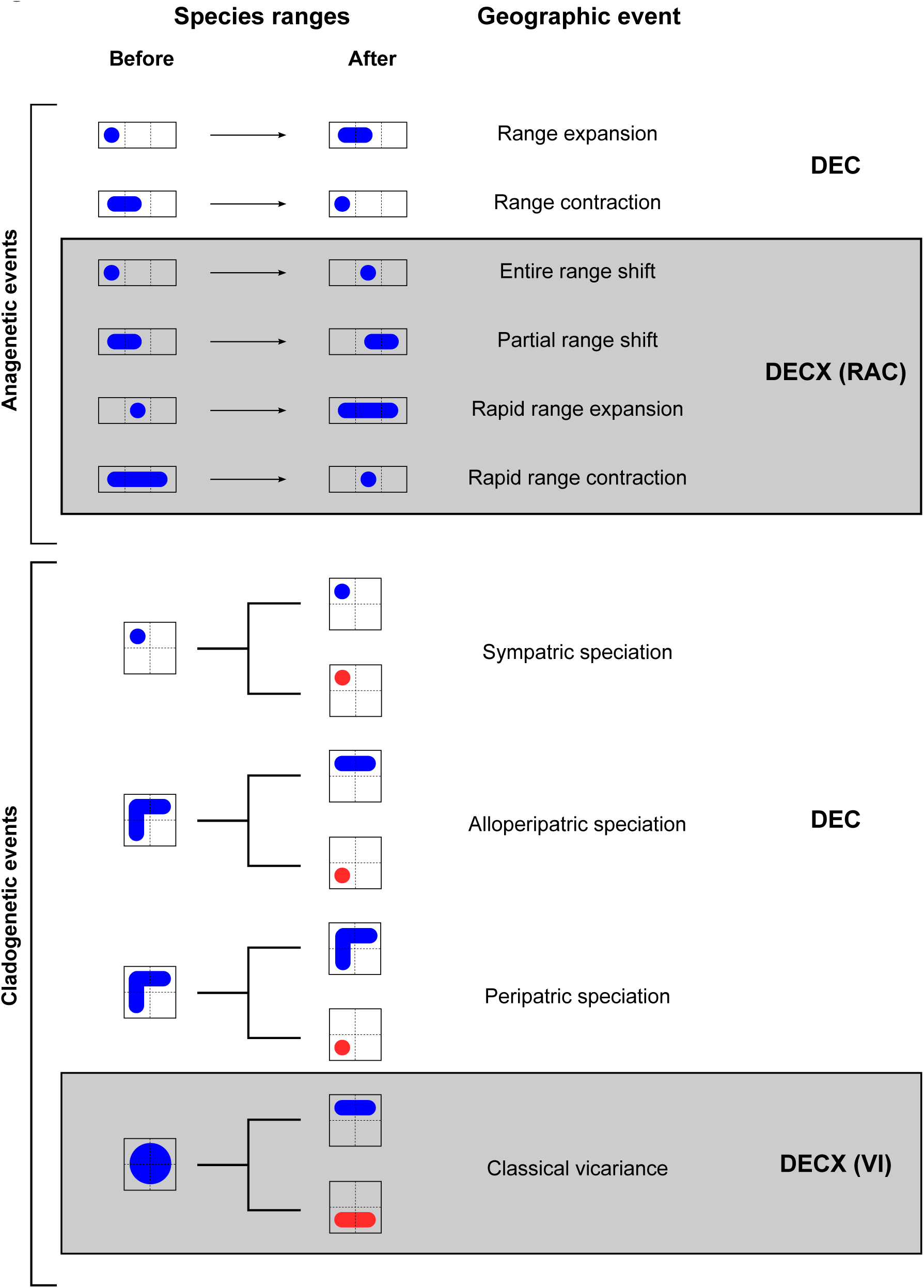
Description of all possible geographic events considered by the *DEC* and *DECX* approaches for modelling range evolution on a phylogeny. Red and blue colours represent species identities, the bold squares/rectangles represent the entire region of study and the dotted lines within delimit land units or areas which can be grouped to form a species range. On a phylogeny, anagenetic events occur along the branches while cladogenetic events correspond to branching events or speciation.

At speciation events or cladogenesis, when an ancestor has a *widely distributed* range (*i.e*. more than a single area), daughter lineages inherit ranges thorough alloperipatry (see Fig. 4a in Ronquist and Sanmartín 2011) or classic peripatry while *single area ranges* are inherited identically (sympatry). In *DECX*, we further add the possibility of classic allopatry or vicariance (as currently implemented in *DIVA*, Ronquist 1997), which means that *both* daughter lineages can inherit ranges formed of more than a single area (*VI*, see **Fig. 2**). For the interested reader, note that this mode of range inheritance is not part of the *DEC+J* model, available as part of the *R*-package *BioGeoBEARS* and as presented by Matzke (2014).

It is also important to note that *DECX* (as in *DEC*) does not support sympatric speciation across multiple areas in which daughter ranges *copy* the entire range of a widely distributed ancestor (as implemented in the *BayArea* approach). Such large scale sympatric speciation does not seem to be a biologically valid assumption on a macroevolutionary scale but which may be applicable to more recent speciation events (Coyne and Orr 2004; e.g. wide sympatric speciation between *Colias eurytheme* and *C. philodice* in North America Wahlberg et al. 2014, or sympatric speciation within several species of *Heliconius* in South America, Kozak et al. 2015). The interested reader can also refer to Ree et al. (2005, p. 2302) for further discussion in this regard.

### Evolving geological connectivity (EGC)

The dispersal rates in *DEC* are generally in the form of an user-specified matrix **D** with its elements representing the effective rate of dispersal from one land unit to another (Ree and Smith 2008). However, some *DEC* based variants (e.g. *SHIBA* or *BayArea*) have opted for a more spatially explicit approach where the area to area dispersal probabilities are a decreasing function of the physical distance separating these areas. Currently, there is no consensus approach other than trying to incorporate as faithfully as possible *a priori* knowledge derived from literature of the various factors influencing the dispersal and establishment of individuals between any two areas. While the **D** matrix is usually symmetric it can also account for asymmetric dispersal due to prevailing winds or ocean currents.

Inference of ancestral states with dispersal specified only by the **D** matrix has its drawbacks as it can produce large ranges with potentially disconnected sub-ranges when the total number of land units is large (Clark et al. 2008). A hypothetical case of spatially disconnected ranges has been illustrated above (see Introduction) in the case of alloperipatric speciation (**Fig. 2**). Moreover, as noted by Buerki et al. (2011, p. 546), when constructing complex biogeographic scenarios using only the **D** matrix, *dispersal over imposed barriers* can still occur due to the non intuitive irreducibility of the **Q** matrix of the CTMC. A simple solution, originally suggested by Ree and Smith (2008), consists of imposing geographical constraints representing between area connectivities such that species ranges are only composed of those areas connected by a migration corridor. These *a priori* constraints are easily viewed in the form of an Undirected Acyclic Graph where the nodes are land units and the vertices represent the possible EGC (**Fig. 3**). The Undirected Acyclic Graph can also be specified as a matrix of binary values in which case it is referred to as an adjacency matrix (Sedgewick and Wayne 2011).

**Figure 3.**
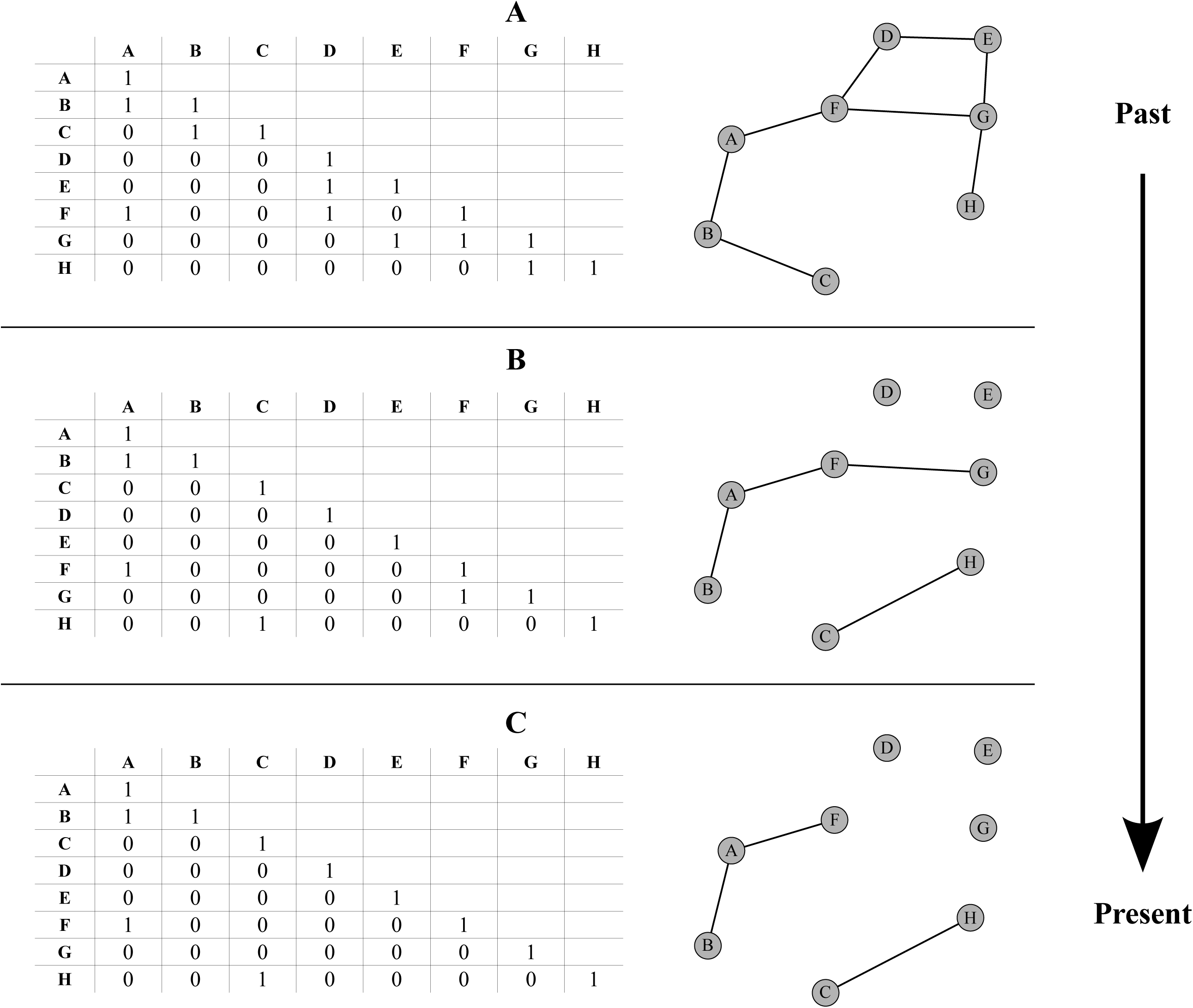
Illustration of an evolving geological connectivity (EGC) over three time intervals (A, B and C). For a given interval, the spatial configuration between areas is defined by the nodes of the graph representing the land units or areas and the lines between them represent their connectivities. Each time interval can also be fully defined by an adjacency matrix composed of zeros and ones respectively specifying the absence or presence of connectivity between any two given nodes.

However, Ree and Smith (2008) and ensuing implementations have only accounted for a single adjacency constraint spanning the entire time scale of the phylogeny. As a result, the majority of studies having used *DEC* or its variants (except *DEC+J*) assume a *static paleo-connectivity* (e.g. Clarke et al. 2008; Clayton et al. 2009; Buerki et al. 2011; Mao et al. 2012; Chacón and Renner 2014). In this paper we attempt to remedy this by implementing an EGC in a computationally efficient manner for the *DEC* and *DECX* models. Conceptually, we make use of a *piecewise time-homogeneous* CTMC framework where the number of Markov states for a given time interval (or *piece*) is conditioned by the respective adjacency matrix. In addition to solving problems related to the irreducibility of the Markov transition matrix, as outlined by Buerki et al. (2011, p. 546), our method is computationally more efficient than multiplying transition probability (**P**) matrices over all the user defined time intervals. Also, our contribution improves upon a crucial and much ignored aspect of the *DEC* model, as noted by Chacón and Renner (2014) that the adjacency matrix strongly determines the outcome of historical inferences in comparison to the dispersal rate matrix **D** even when the latter is allowed to vary across time intervals.

### PODs: biogeographic context

We define our hypothetical time-stratified paleogeographic scenario in which the EGC crudely imitates the breakup and drift of Pangea up to our main present day configuration of land masses. This scenario consists of three time intervals starting with eight well connected land units (**Fig. 3A**) in the past which progressively break away into smaller units or regroup into different ones (**Fig. 3B**) and finally give rise to a present day configuration (**Fig. 3C**). Each interval in time thus consists of a corresponding adjacency matrix, a dispersal rate matrix which specifies the amplitude and directionality of dispersal (*i.e*. A→B and B→A) between land units.

### PODs: phylogenies used

Time-calibrated trees were generated under various *birth-death* scenarios (e.g. expanding diversity, declining diversity, high turnover) which represent the most commonly found patterns of lineage diversification in the literature. We used the *R*-package *TESS* (v.2.0, Höhna 2014) and the function *sim.globalBiDe.age* for simulating the time-calibrated ultrametric phylogenies under a global time dependent birth-death process conditioned on the age of the tree. The latter was set to correspond to the sum of the three equally spaced time intervals. Thus, lineage diversification during these three intervals, each lasting over 10 time units, consisted of *piecewise constant* speciation and extinction rates. Totally, 12 diversification scenarios were considerd and more details on each of these can be found in the Dryad repository (Online **Appendix 1** available at https://goo.gl/oXDdJz).

For our PODs study, we chose not to generate trees under any of the State-dependent Speciation and Extinction models (*SSE*, Maddison et al. 2007; Goldberg et al. 2011) for several reasons. First, large phylogenies (300+ species) are needed to get a significant signal between rates of speciation/extinction/transition and a given trait (Davis et al. 2013), and we also wanted to illustrate the performance of *DECX* on small phylogenies. Second, it quickly becomes intractable to handle large phylogenies with multiple geographic states (our simulation study area consists of eight areas). Third, while the *GeoSSE* model (Goldberg et al. 2011) can account for a geographic state composed of more than one area, it still allows only a maximum of two areas which would be impossible to accommodate in our biogeographic set up with eight areas. Fourth, the mathematical foundations of *SSE* approaches (Maddison and FitzJohn 2015) have recently been called into question based on concern from false positive results when using these models (Rabosky and Goldberg 2015).

### PODs: analyses

Given a phylogeny, dispersal rate matrices (**D**) and an EGC, biogeographic histories are simulated forwards in time which result in hypothetical extant range distributions in terms of species presence-absences. Two baseline models, *DEC* and *DEC-VI+RAC*, were chosen to generate the PODs while inference was performed using the following models *DEC*, *DEC-VI, DEC-RAC* and *DEC-VI+RAC* (**Fig. 2**) *i.e.* with either the true model or a variant of the latter. For a given baseline model and lineage diversification scenario, 5000 PODs were generated each with a randomly chosen dispersal (*d*) and extinction (*e*) parameter in the interval [0, 0.2] and with *e < d* in order to avoid highly trivial scenarios of extant species ranges. Note that we exclude CTMC state changes towards the null range in our PODs and our inference as we are dealing only with ultrametric trees. This gave us a total of 2 (true models) x 4 (inference models) x 12 (diversification scenarios) x 5000 = 480,000 biogeographic inferences to be analyzed. Each inference thus consists of estimates for the *d* and *e* parameters along with the entire biogeographic history *i.e.* the inferred ancestral ranges at each interior node of the respective phylogeny.

### Empirical analysis of island clades (Matzke 2014)

The *DECX* model was not specifically developed for studying either island or continental biogeography and our search for recently published and freely available studies using *DEC* and alternate models on real data yielded Matzke (2014, Table 2). The latter contains a compilation of 13 island clades (mostly from the Hawaiian archipelago) for which the biogeographic histories have been inferred under a variety of hypotheses and where model choice was evaluated between the *DEC* and *DEC+J* approaches. We made use of these easily accessible datasets (Matzke 2014, Dryad: http://dx.doi.org/10.5061/dryad.2mclt) as it sets a good precedent for the reproducibility of results in the field of historical biogeography. Note that some of the scenarios in Matzke (2014, Table 2) specify a *ghost source* or an “ancient area *Z*” representing either an island older than the current Hawaiian Islands or the continent. For these cases we have modified the adjacency matrix such that Z connects at least once with the first of the emerging islands *i.e.* Kauai (or “*K*”), Without this modification, our implementation returned infinite likelihoods. This situation can be understood as follows: in the most ancient time interval, *Z* remains the sole possible landmass but stays disconnected with all other present day areas and throughout all other time intervals. Thus, a jump dispersal event remains the sole compatible and crucial event for calculating the likelihoods at the root node. For each island clade in the original study (Matzke 2014, Table 2), various biogeographic hypotheses are presented for which we compare results obtained under our implementation of the *DEC* (hereafter C++ *DEC*) and *DECX* models to the already published *BioGeoBEARS-DEC* and *DEC+J* results. However, a notable difference from Matzke (2014, Table 2 and Supplementary Table 1) is that out of a total of 53 biogeographic analyses we have selected only 44 (*cf. Difficulties with the empirical analysis of island clades*). Note that in this following we shall refer to *DECX* instead of specifying the *DEC-VI+RAC* sub-model to be consistent with the fact that we haven’t explored the *VI* and *RAC* aspects separately on the empirical analyses (as has been done for the PODs analysis).

**Table 2.** The average proportion of accurately inferred ancestral ranges for each of the 12 diversification scenarios defined in **Appendix 1** (available online at https://goo.gl/oXDdJz) and for each true model/inferred model combination. Global averages over 12 scenarios are also provided.

| Simulation model | Inference model | Global | Scenario 1 | Scenario 2 | Scenario 3 | Scenario 4 | Scenario 5 | Scenario 6 | Scenario 7 | Scenario 8 | Scenario 9 | Scenario 10 | Scenario 11 | Scenario 12 |
| --- | --- | --- | --- | --- | --- | --- | --- | --- | --- | --- | --- | --- | --- | --- |
| DEC | DEC | 0.94 | 0.97 | 0.96 | 0.93 | 0.98 | 0.91 | 0.94 | 0.96 | 0.9 | 0.97 | 0.92 | 0.93 | 0.9 |
|  | DEC-VI | 0.94 | 0.96 | 0.96 | 0.93 | 0.98 | 0.91 | 0.94 | 0.96 | 0.9 | 0.97 | 0.92 | 0.93 | 0.9 |
|  | DEC-RAC | 0.91 | 0.93 | 0.93 | 0.88 | 0.96 | 0.87 | 0.91 | 0.94 | 0.86 | 0.95 | 0.9 | 0.89 | 0.87 |
|  | DEC-VI+RAC | 0.91 | 0.93 | 0.93 | 0.88 | 0.96 | 0.87 | 0.91 | 0.94 | 0.86 | 0.95 | 0.89 | 0.89 | 0.87 |
| DEC-VI+RAC | DEC | 0.82 | 0.89 | 0.84 | 0.7 | 0.94 | 0.73 | 0.85 | 0.89 | 0.74 | 0.93 | 0.82 | 0.81 | 0.75 |
|  | DEC-VI | 0.83 | 0.89 | 0.85 | 0.7 | 0.94 | 0.74 | 0.86 | 0.89 | 0.74 | 0.93 | 0.82 | 0.82 | 0.76 |
|  | DEC-RAC | 0.85 | 0.91 | 0.87 | 0.73 | 0.95 | 0.77 | 0.89 | 0.92 | 0.78 | 0.95 | 0.84 | 0.84 | 0.77 |
|  | DEC-VI+RAC | 0.86 | 0.92 | 0.87 | 0.74 | 0.96 | 0.78 | 0.89 | 0.93 | 0.79 | 0.95 | 0.85 | 0.85 | 0.78 |

### Empirical analysis of Amphibia (Pyron 2014)

In order to further illustrate the efficiency and performance of the *DECX* approach when applied to large phylogenies, we chose an Amphibian phylogeny containing 3309 species (Pyron 2014). We also make use of the species distribution and area delineation as presented by Pyron (2014) who applied the *BioGeoBEARS-DEC* and *DEC+J* models to the Amphibian dataset. In the following, and only for the Amphibian dataset, we define as model **M0** whenever inference was performed without imposing any EGC. Note that we constrain our **M0**, akin to Pyron (2014), such that the inferred ancestral ranges can only be composed of a maximum of two areas. We first ran the *BioGeoBEARS-DEC* and *DEC+J* models using the input files provided by Pyron (2014, Dryad: http://dx.doi.org/10.5061/drvad.im453) and adapted the latter for the C++ *DEC* and *DECX* analysis. Next, we imposed an EGC covering the lineage diversification history of Amphibians from the present to the past 270 Myrs which we shall refer to as model **M1**. The EGC for the Amphibian history was thus divided into five time intervals: (*i*) 0 to 34 Myrs ago, (*ii*) 34 to 90 Myrs ago, (*iii*) 90 to 120 Myrs ago, (*iv*) 120 to 170 Myrs ago, and (*v*) 170 to 270 Myrs ago. Note that the “maximum of two areas” constraint applies to the **M1** model as well. As the latter constraint might seem unreasonable, especially for ancient clades which may have had a geographic spread over wide continental landmasses (e.g. Pangea, Gondwana, Laurasia), we extend the **M1** model to have a maximum number of areas per ancestral range to four, which will be referred to as the **M2** model.

### Empirical analysis of palpimanoids (Wood et al. 2013)

Trees with fossil tips (*i.e*. non-ultrametric) can now be analyzed explicitly by the *DECX* framework. Previous implementations of *DEC* (especially the C++ version as available at https://github.com/blackrim/lagrange) accept trees with fossil tips without complaint and erroneously extend the respective branches up to the present. This is due to an internal procedure for calculating the phylogenetic branch lengths from node age information provided by the user in the NEWICK format. In the *DECX* implementation, we have corrected this by asking the user to explicitly specify when providing a non-ultrametric tree and accordingly account for the correct branch length of the fossil tips. We compare our implementation by reanalyzing the biogeography of palpimanoid spiders (Wood et al. 2013; Dryad: http://dx.doi.org/10.5061/drvad.7231d.2) which made use of the *DEC* model without the fossil tip correction. Their phylogeny comprises 23 lineages of which five are fossil tips.

### Implementation

The *DECX* approach for inferring dispersal and extinction parameters along with the ancestral species ranges is essentially coded in C++ and can be freely downloaded from the GitHub repository https://github.com/champost/DECX. This is an overhaul of the original C++ version of the *DEC* model (Smith 2009; https://github.com/blackrim/lagrange). Among some of the notable improvements, it is now possible to analyze large phylogenies (over 1000 species) which previously resulted in bugs related to the high precision required for the calculation of the likelihood. Another area of interest is the handling of a few highly dispersed (widespread) extant species of a given clade, which is usually an artifact of recent and rapid expansion and which can now be modelled separately. The *DECX* program is meant to be used from the command line and simply requires a single configuration file and several input files (e.g. trees in NEWICK format). Additional information can be found at the aforementioned website.

## Results

### PODs

Overall model performance of the approaches introduced in this paper (*i.e. DEC, DEC-VI, DEC-RAC*, and *DEC-VI+RAC*) with respect to one another was evaluated using a pairwise comparison of the respective maximum likelihoods (**Fig. 4;** Online **Appendix 2** available at https://goo.gl/oXDdJz). Note that we adopted this strategy as all these models have the same number of degrees of freedom which excludes likelihood ratio tests and all things being equal in our case, the use of information theoretic measures such as Akaike Information Criterion basically amounts to a comparison of the model likelihoods. When *DEC* is the true underlying model of range evolution **Fig. 4** (top row) it always has a lower or equal *-InL* than the other three models. In this case, the *DEC-VI* model agrees almost perfectly with the *DEC* inferences and note that the other two models (*DEC-RAC* and *DEC-VI+RAC*) disagree in a similar fashion. When *DEC-VI+RAC* is the true model **Fig. 4** (bottom row) *DEC-RAC* inferences are in very good agreement with the former compared to the other two models (*i.e. DEC* and *DEC-VI*). Additional results illustrating the effect of the phylogeny, depending on the various lineage diversification scenarios, on model comparison using likelihoods can be found in the Dryad repository (Online **Appendix 3** available at https://goo.gl/oXDdJz).

**Figure 4.**
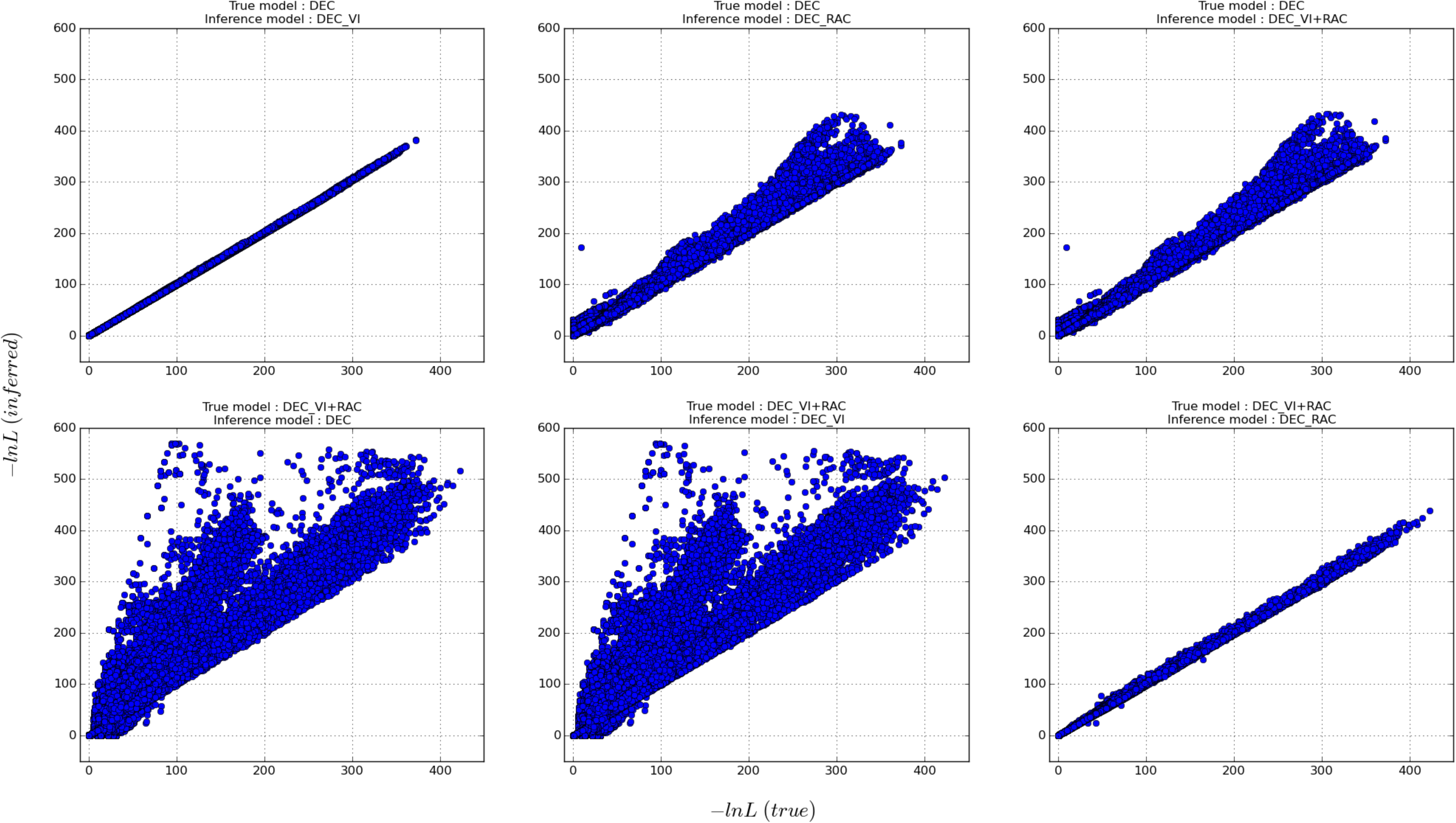
Comparison of the inferred -*ln*L values on Pseudo Observed Datasets (PODs). The true model is the one that generated the PODs and the inferred model is every other alternate model. The true model for the top and bottom rows are *DEC* and *DEC-VI+RAC* respectively. The complete results are available on Dryad (Online **Appendix 2** available at https://goo.gl/oXDdJz).

Likelihoods aside, performance was also evaluated by a relative comparison of the inferred dispersal (*d*) and extinction (*e*) parameters with their true values. **Fig. 5** plots various true model versus inferred model scenarios for which relative proportions of inferred dispersal and extinction are shown (see also Online **Appendix 4** available at https://goo.gl/oXDdJz). Overall, both parameters are underestimated and seem to follow a general hyperbolic relationship where the performance of one of the parameter relates inversely to the other. However, irrespective of the true model, inference under *DEC-RAC* and *DEC-VI+RAC* introduces more bias in dispersal estimates in comparison to the other two models. Likewise, some estimates of extinction are more biased when using the *DEC* and *DEC-VI* models.

**Figure 5.**
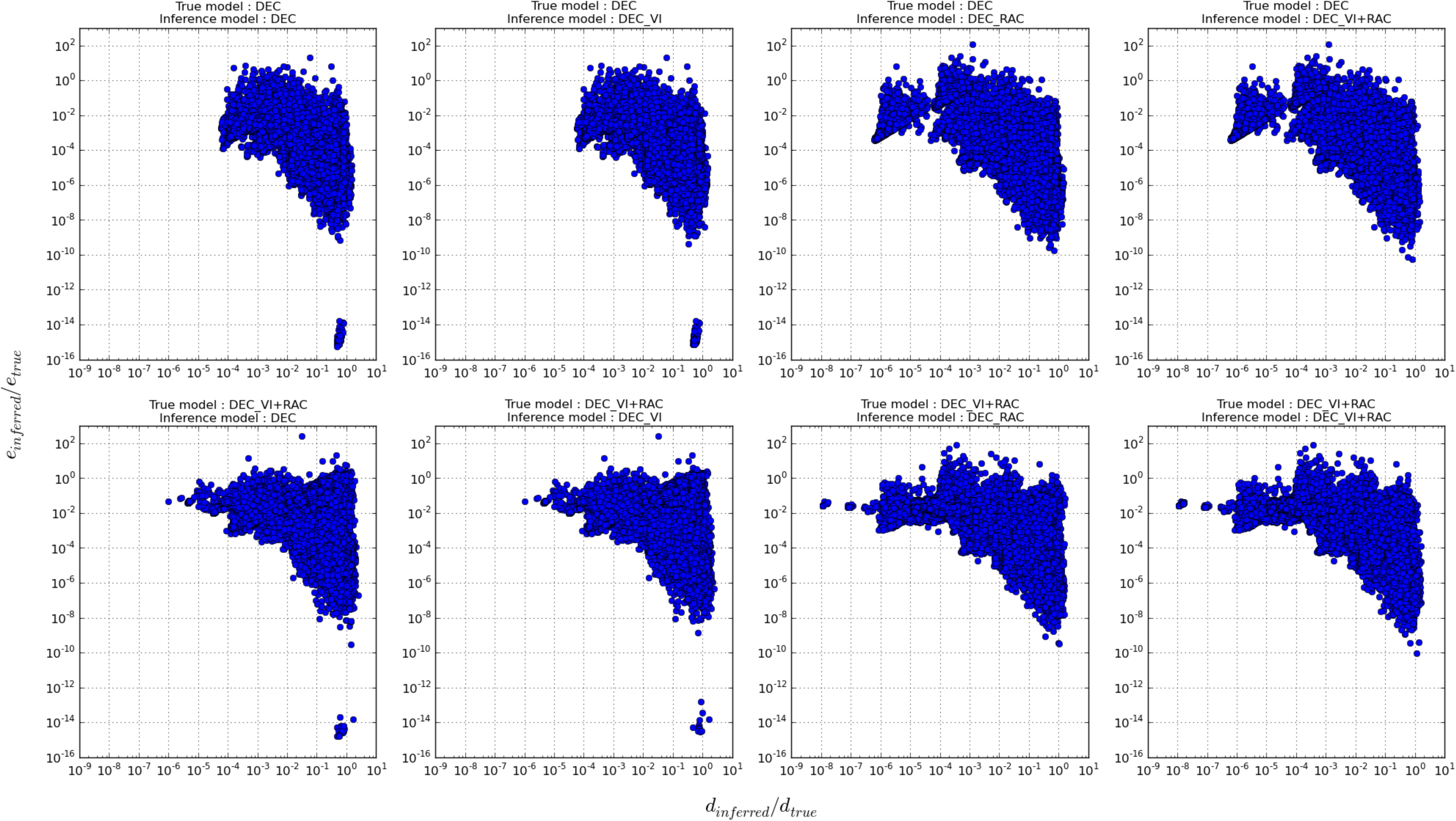
Scaled parameter inferences (with respect to their true simulated values) of the dispersal rate (*d*) plotted against the local extinction rate (*e*) for different model assumptions. All points that lie on *y* = 10° = 1 are unbiased inferences of the true local extinction rate and those that lie on *x* = 10° = 1 are unbiased inferences of the true dispersal rate. The complete results are available on Dryad (Online **Appendix 4** available at https://goo.gl/oXDdJz).

Model performance was also evaluated in terms of the proportion of accurately inferred ranges. Globally, over approximately half a million inferred histories, the true underlying models are also consistently the models with the highest proportion of accurately inferred ranges (**Table 2**). With *DEC* as the true model of range evolution, all models are above 90% and the proportions concur with their respective differences in model likelihoods. However, when *DEC-VI+RAC* is the true model, all inference models are less accurate (above 80%) and the proportions between models differ but slightly. We also explored the effect of the number of simulated ranges and the size of the phylogeny (*i.e*. number of tips) on the proportion of accurately inferred ranges (Online **Appendix 3** available at https://goo.gl/oXDdJz). Apart from a general trend where an increasing number of simulated ranges decreases the proportion of accurately inferred ranges, we did not find anything conclusive.

### Empirical analysis of island clades (Matzke 2014)

**Table 3** of this paper presents our biogeographic analysis of 13 island clades, mainly from the Hawaiian archipelago (Matzke 2014, Table 2). Alternative model constraints on these clades and subsequent difficulties (*cf. Difficulties with the empirical analysis of island clades*) result in a total of 44 analyses for which we report the -*ln*L values obtained under the C++ *DEC* and *DECX* models alongside those already available for *BioGeoBEARS-DEC* and *DEC+J*. Previously, Matzke (2014, Fig. 5) indicated that 51/53 island clades analyses support, in terms of AICc model weights, the *DEC+J* model over the *BioGeoBEARS-DEC* model. Our study finds a more nuanced pattern depending on the model assumptions and subsequent constraints. Out of the 44 considered analyses (**Table 3**), we find that 48% (*i.e*. 21/44) were better explained by the *DECX* model whereas 52% of the analyses (23/44) are in favour of the *DEC+J* model.

**Table 3.**
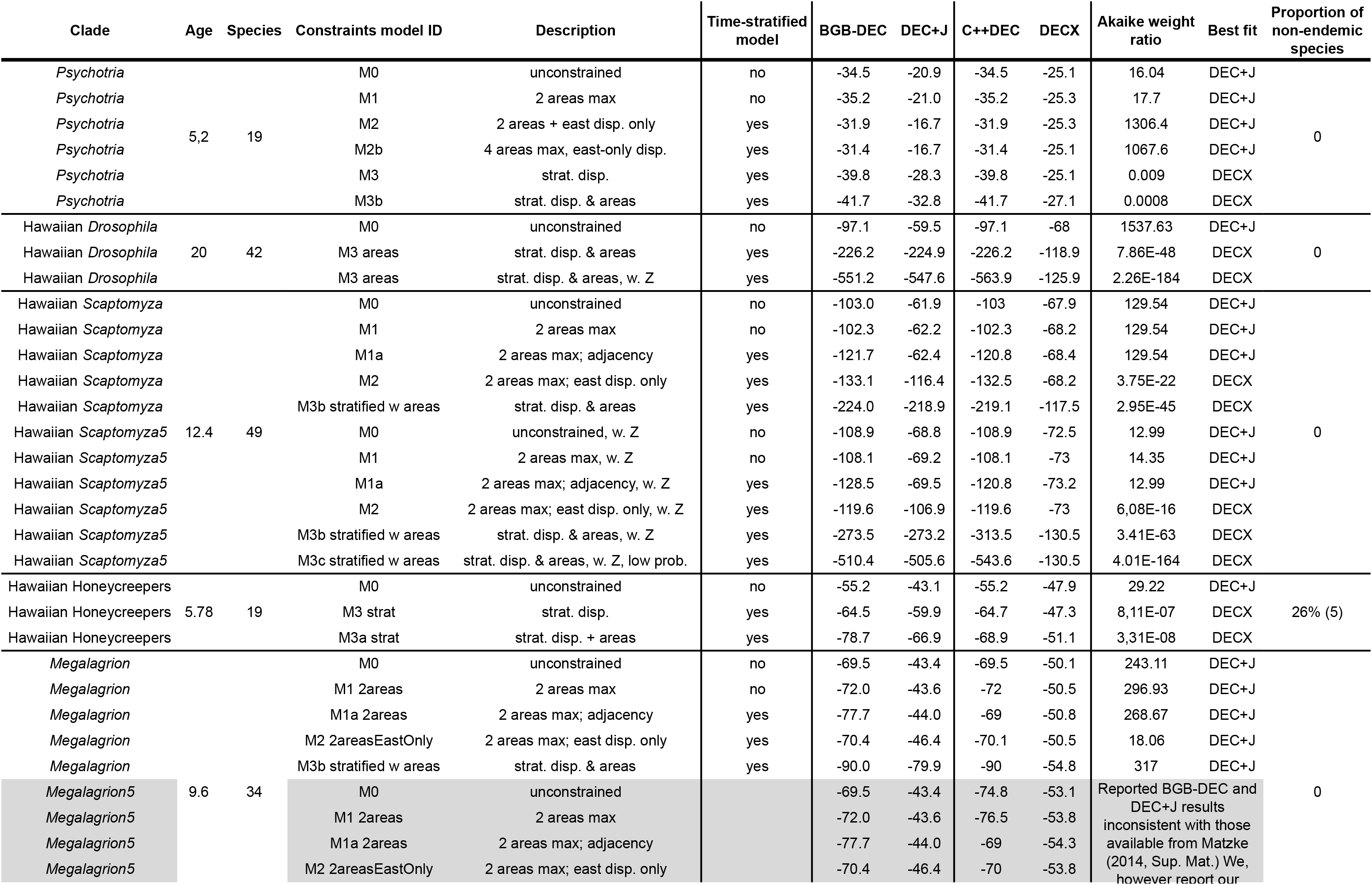
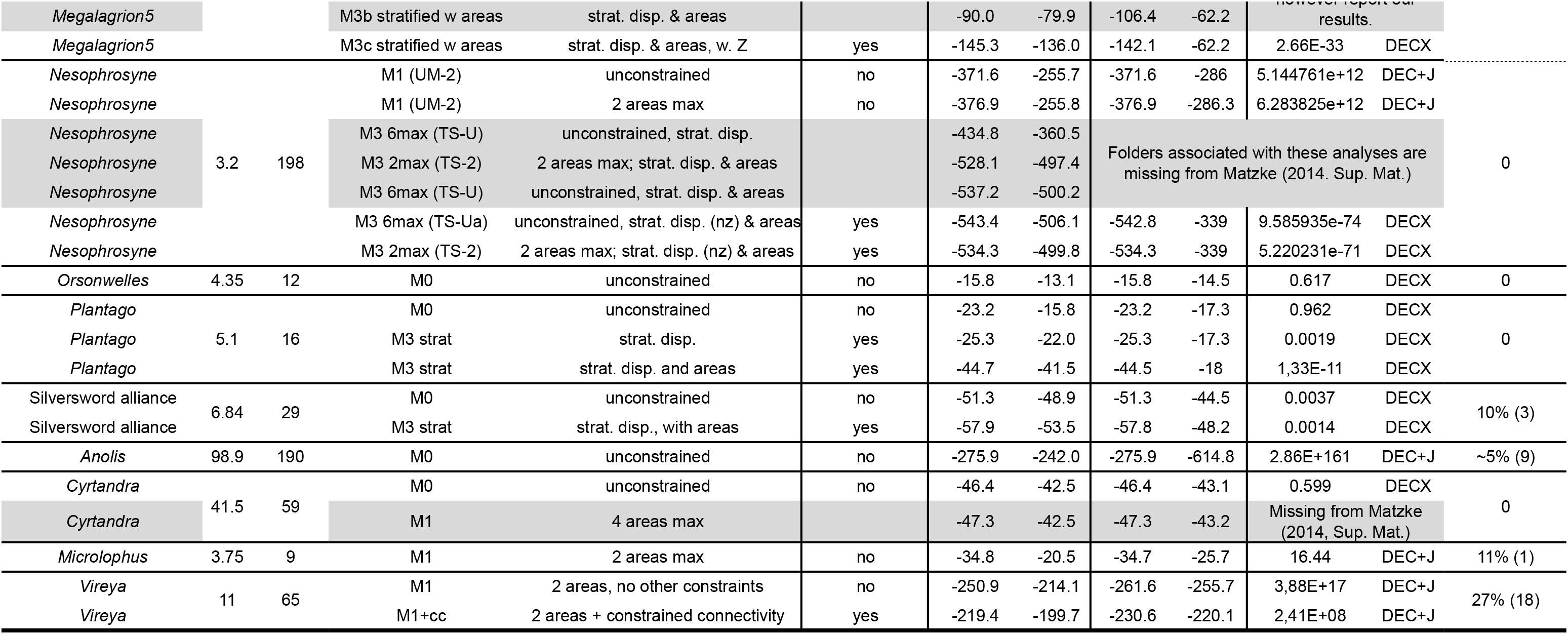
Model performance comparing the *DEC*, *DEC+J*, and *DECX* models on 13 island clades comprising a total of 53 biogeographic scenarios (adapted from Matzke, 2014). AICc and Akaike weight ratio were used for model comparison. Abbreviations: straf, time-stratified; disp., manual dispersal probability multiplier matrix; w. Z, analysis was run with “Z”, an old, ancestral area outside of the extant Hawaiian high islands; nz, zeros in the manual dispersal probability multiplier matrix have been replaced with a small non-zero value (*cf* Matzke, 2014 for further details). For further information on the rows greyed out, refer to *Difficulties with the empirical analysis of island clades* in the main text. The complete results are available on Dryad (Online **Appendix 5** available at https://goo.gl/oXDdJz).

Two types of constraint scenarios particularly highlight the differences in performance between the *DECX* and *DEC+J* models (**Table 3**). They are no constraints or constraints based either on the maximal size of the inferred ancestral ranges (referred to here as *weakly constrained*) and those that are time-stratified (sometimes combined with the former). Out of the 19 weakly constrained analyses, we find that 21% (*i.e*. 4/19) were better explained by the *DECX* model whereas 79% of the analyses (15/19) were in favour of the *DEC+J* model. Out of the 25 time-stratified analyses we find that 68% (*i.e*. 17/25) are better explained by the *DECX* model whereas 32% of the analyses (8/25) were in favour of the *DEC+J* model (*cf. Inferring geographic range evolution using DECX and DEC+J*). Note that these proportions provide only a *qualitative insight* into the performance of the *DECX* model versus the *DEC+J* model. While Matzke (2014, pg. 956) states that, “*Wherever a constrained analysis was run, an unconstrained run was also implemented for comparison*.”, we note that this hasn’t been followed rigorously. The most obvious example being the 100 Myr old Anolis clade in the Caribbean where only a single unconstrained analysis has been performed. Following Matzke (2014), we provide the input files necessary to redo our analyses, as well as additional information such as the trees displaying the inferred ancestral areas and the raw program output (*i.e*. with -*ln*L values, list of ranges within 2 *ln*L units for every ancestral node, etc.) in the Dryad repository (Online **Appendix 5** available at https://goo.gl/oXDdJz).

### Empirical analysis of Amphibia (Pyron 2014)

For the **M0** model (*i.e*. no additional constraints) and under *BioGeoBEARS-DEC* we obtained very different values for the -*ln*L and for estimates of dispersal and extinction from those of Pyron (2014), though the inferred origin of the Amphibian phylogeny was consistent with our results (**Table 4**). As expected, C++ *DEC* agrees more closely with the *BioGeoBEARS-DEC* analysis in our paper except for the extinction parameter. We also found a notable difference in -*ln*L values between the *DEC+J* analysis performed by Pyron (2014) and our results which favour the *DEC+J* model as the best-fitting model of the historical biogeography of Amphibians. The *DECX* model fits the Amphibian dataset poorly under the **M0** assumption though the inferred origin and dispersal rate remains consistent with the other models.

**Table 4.** Comparison of the *DEC*, *DEC+J* and *DECX* model performance for the Amphibian phylogeny from Pyron (2014). For each model, the log-likelihood (LnL) value, the number of free parameters (Param.), as well as the parameter estimates for *d* (dispersal), *e* (extinction), and *j* (founder-event speciation) are reported. The ancestral origin of the clade and the associated relative probability are also shown (AF: Africa; NEA: Nearctic; NT: Neotropics; and PA: Palearctic). For more information on the inferred biogeographic history, see Fig. 6. The complete results are available on Dryad (Online **Appendix 6** available at https://goo.gl/oXDdJz).

| Model | Analysis | Description | LnL | Param. | d | e | j | Inferred origin | Rel. Prob. |
| --- | --- | --- | --- | --- | --- | --- | --- | --- | --- |
| BGB-DEC | Pyron (2014) | M0 unconstrained, max. 2 areas | -1721.2 | 2 | 0.000753 | 0.000753 | 0 | NT + NEA | ≈0.63 |
| BGB-DEC | This study | M0 unconstrained, max. 2 areas | -1617.66 | 2 | 0.000356 | 1,00E-12 | 0 | NT + NEA | 0.66 |
| C++DEC | This study | M0 unconstrained, max. 2 areas | -1617.95 | 2 | 0.000368 | 4.726e-06 | 0 | NT + NEA | 0.66 |
| DEC+J | Pyron (2014) | M0 unconstrained, max. 2 areas | -1804.3 | 3 | 0.000996 | 0.000982 | 0.000997 | NT + NEA | ≈0.6 |
| **DEC+J** | **This study** | **M0 unconstrained, max. 2 areas** | **-1550.96** | **3** | **0.000244** | **1,00E-12** | **0.0015** | **NT + NEA** | **0.67** |
| DECX | This study | M0 unconstrained, max. 2 areas | -7899.73 | 2 | 0.000226 | 0.01894 | 0 | NT | 0.03 |
| BGB-DEC | This study | M1 strat. adjacency, max. 2 areas | -1782.92 | 2 | 0.000378 | 0.000588 | 0 | NEA + AF | ≈0.9 |
| C++DEC | This study | M1 strat. adjacency, max. 2 areas | -1670.2 | 2 | 0.001752 | 0.000565 | 0 | NEA + AF | 0.59 |
| DEC+J | This study | M1 strat. adjacency, max. 2 areas | -1571.13 | 3 | 0.000224 | 1E-12 | 0.001961 | NT + NEA | ≈0.9 |
| **DECX** | **This study** | **M1 strat. adjacency, max. 2 areas** | **-1496.5** | **2** | **0.000384** | **0.000116** | **0** | **NT + NEA** | **0.64** |
| BGB-DEC | This study | M2 strat. adjacency, max. 4 areas | -1835.58 | 2 | 0.000311 | 0.001979 | 0 | NEA + AF + PA | ≈0.9 |
| C++DEC | This study | M2 strat. adjacency, max. 4 areas | -1587.9 | 2 | 0.001483 | 0.000509 | 0 | NT + NEA + AF + PA | 0.14 |
| DEC+J | This study | M2 strat. adjacency, max. 4 areas | -1572.84 | 3 | 0.000212 | 1E-12 | 0.0019297 | NEA + AF + PA | ≈0.8 |
| **DECX** | **This study** | **M2 strat. adjacency, max. 4 areas** | **-1515.6** | **2** | **0.000348** | **6,00E-05** | **0** | **NT + NEA + PA** | **0.3** |

When we define an EGC through time (models **M1** and **M2**), *DECX* fits the data considerably better with respect to the other models. It is also worth noting that under the **M1** model the -*ln*L values for C++ *DEC* and *BioGeoBEARS-DEC* differ even though the inferred parameters (especially *e*) and the species range at the tree origin are in agreement. This difference is greater for the **M2** model where the inferred parameters and ancestral ranges also differ. For both **M0** and **M1** and with a lesser number of parameters, *DECX* turns out to be the more plausible fit to the data with respect to the *DEC+J* model. Other than the -*ln*L and the extinction parameter, *DECX* and *DEC+J* are consistent in terms of the inferred dispersal and the range at the origin of the phylogeny.

We also attempted a more detailed comparison of the entire biogeographic history inferred by the *DEC+J* and *DECX* models which differed mainly at the deeper nodes (**Fig. 6**). The ancestor of all amphibians is found to be in a range composed of Neotropics, Nearctic, and Palearctic with *DECX*, whereas is found to be in Africa, Nearctic, and Palearctic with *DEC+J* (the same ranges are found for the ancestor of the clade Anura + Caudata). *DECX* infers an ancient vicariance event splitting the ancestor of the clade Anura + Caudata, which is the result of Anura splitting in the Neotropics and Caudata splitting in Nearctic and Palearctic. Within Caudata, both methods infer a similar (if not identical) biogeographic pattern. However, within Anura, both methods differ slightly. *DECX* estimates that the Neotropics were the ancestral region of the deepest nodes, while *DEC+J* infers the range composed of Africa, Nearctic, and Palearctic (or Neotropics, Nearctic, and Palearctic). Other important differences at particular nodes are highlighted within the dashed boxes (**Fig. 6**). *DECX* finds an ancestral range composed of the Neotropics, Africa, and India to be split into Neotropics for one clade, and Africa plus India for the second clade. On the contrary, *DEC+J* finds an ancestral range composed of the Neotropics, Africa, and Australasia to be split into Neotropics for one clade, and Africa plus Australasia for the second clade. All figures displaying the inferred ancestral ranges as well as the files with the raw results (*i.e. -ln*L, list of ancestral ranges within 2-log *ln*L units, etc.) are available in the Dryad repository (Online **Appendix 6** available at https://goo.gl/oXDdJz).

**Figure 6.**
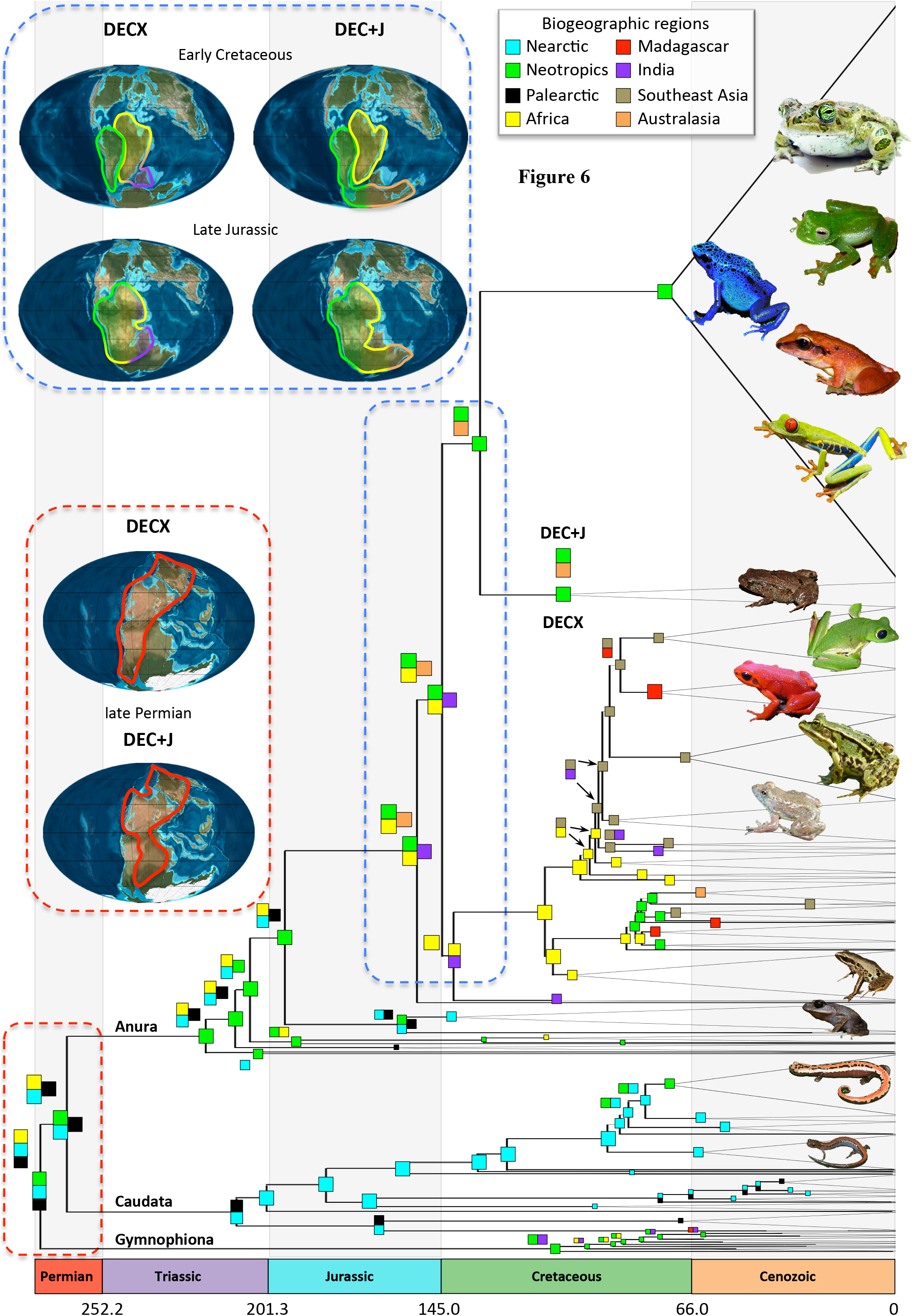
Comparison of the biogeographic histories as inferred by the *DECX* and *DEC+J* models for the Amphibians. The best-fit biogeographic model under *DECX* (see also **Table 4**) corresponds to the M2 model (is plotted) which includes a time-stratified paleogeographic model and restricts the ancestral range size to maximum four areas. *DECX* results are depicted at nodes, and *DIX+1* results (when different from *DECX*) are depicted just above. Important discrepancies are highlighted by dashed boxes, which are also portrayed on paleogeographic maps (Blakey, 2008). The complete results are available on Dryad (Online **Appendix 6** available at https://goo.gl/oXDdJz).

### Empirical analysis of palpimanoids (Wood et al. 2013)

Our results of the biogeography of palpimanoid spiders highlighted the fact that fossils, when used in a total evidence framework, should have a strong influence close of their respective branching points. We thus identified five ancestral nodes (**Fig. 7**), in the vicinity of the fossil tips, of the palpimanoid phylogeny which differ from the previous analyses by Wood et al. (2013). In terms of parameter estimates, Wood et al. (2013) found a smaller extinction rate with respect to dispersal (*d*=5.36×10^-4^ and *e*=6.85×10^-5^), Though, we find an extinction rate smaller than dispersal, we also obtain a more than two-fold increase in the former (*d*=6.42×10^-4^ and *e*=1.62×10^-4^). The *DECX* analysis with the same dataset and same parameters resulted in an extinction rate almost twice as high as dispersal rate (*d*=2.71×10^-4^ and *e*=4.36×10^-4^). All analyses and detailed results are available in the Dryad repository (Online **Appendix 7** available at https://goo.gl/oXDdJz).

**Figure 7.**
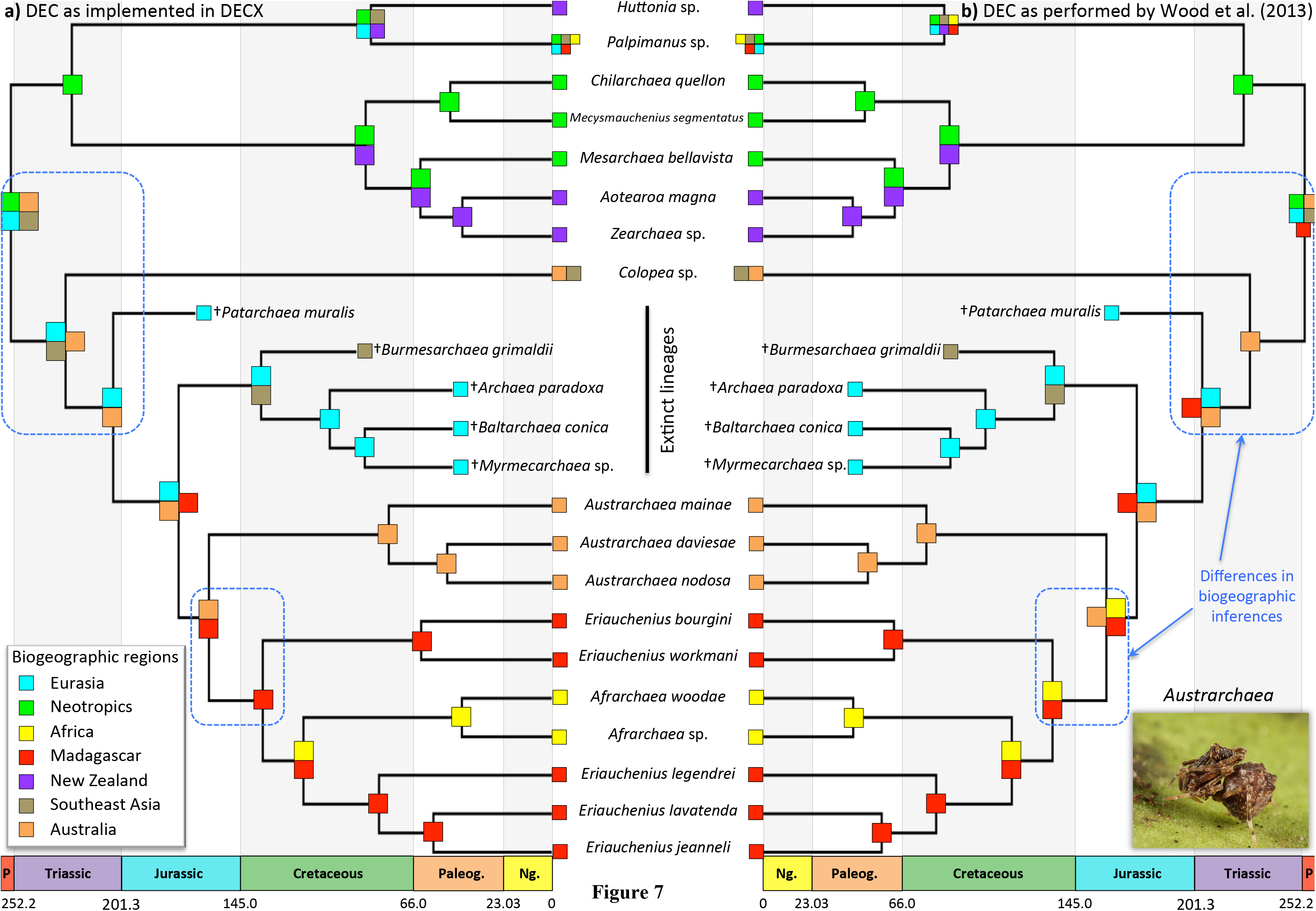
Comparison of biogeographic histories using trees with fossil tips for palpimanoid spiders (Wood et al. 2013) as inferred by a) C++ *DEC* (as implemented in *DECX*) and b) *DEC* as available from https://github.com/blackrim/lagrange. The former accounts for non-ultrametric trees while the latter artificially extends fossil tips to the present. Fossil lineages are those branches stopping before reaching the present and highlighted by a cross sign. Important discrepancies in biogeographic inferences between the two models are highlighted by dashed boxes. The complete results are available on Dryad (Online **Appendix 7** available at https://goo.gl/oXDdJz).

## Discussion

### Model assumptions in parametric biogeography

Currently, a plethora of parametric models are available for the study of historical biogeography (Ree and Sanmartín 2009; Ronquist and Sanmartín 2011; Wen et al. 2013; Matzke 2014). By developing the *DECX* framework in this paper, we have added yet another conceptual and methodological avenue which makes use of phylogenies to study biogeography. This begs the question: how does *DECX* differ from previous approaches ? In the following, we review some of the salient model assumptions as a guideline for the interested reader and a potential user.

#### DIVA

As a precursor to process-based biogeographic models, *DIVA* (Ronquist 1997) estimates ancestral ranges in a parsimonious framework based on minimizing costs associated with events. Speciation occurs either allopatric, through the vicariance of an ancestral range into two disjoint subsets, or by sympatry within a single area. *DIVA’*s parsimony framework dictates that cladogenetic events have a zero cost whereas dispersal and extinction events come at an increased cost proportional to the number of concerned areas. The simplicity of its assumptions results in vicariance being favoured when data are scarce and a profusion of dispersal events at the tips of the phylogeny (Kodandaramaiah 2010; Buerki et al. 2011). While *DIVA* ignores branch length information and neither is it capable of using an EGC to inform its inference of ancestral areas, recent additions now make it possible to integrate over topological uncertainty (Nylander et al. 2008). A notable achievement of the *DECX* framework has thus been to integrate *DIVA* style vicariant mode of speciation, previously absent in any of the *DEC* based approaches (see Fig. 4a in Ronquist and Sanmartín 2011).

#### DEC

As expounded in the previous sections, *DECX* maintains the spirit of the *DEC* approach with regards to model complexity as represented by the number of free parameters. **Fig. 2**. further elaborates on the differences in the anagenetic and cladogenetic processes accounted by both the methods. In summary, the original *DEC* model (Ree and Smith, 2008) remains a special case of *DECX* (*cf. Inferring geographic range evolution using DEC and DECX*).

#### BayArea

This is very first approach which accommodates for fine-scale biogeographic processes (*i.e*. a study system with a large number of areas) and in a computationally efficient manner (Landis et al. 2013). *BayArea* also takes the cue from Webb and Ree (2012) to model anagenesis as affected by extrinsic factors such as distance. Though, these capabilities come at the price of highly restrictive model of geographic speciation (sympatry irrespective of the ancestral range size) and a static landmass configuration throughout the timescale of the phylogeny. While, *BayArea* can be extended to include many of the features of *DEC/DECX*, a lot of this progress would hinge upon a particularly tricky specification of the Metropolis-Hastings ratios which lie at the heart of the MCMC exploration of biogeographic histories. In comparison to *BayArea*, the use of a *piecewise time-homogeneous* CTMC approach greatly helps the *DECX* model to also handle a large number of areas. Though, the precise nature of this advantage will depend on the biological problem at hand and the corresponding EGC framework which is an open topic for future study (*cf. Advantages of the DECX framework*).

#### SHIBA

Very simply put, *SHIBA* can be viewed as a forwards-in-time version of the the *DEC* model with the ML inference step replaced by basic rejection sampling. *SHIBA* is also one of the first models which incorporates the inference of local extinction rates based on the size of the areas and a **D** matrix with dispersal rates scaled as a function of distance. While a forwards-in-time version implementation is one of the most flexible frameworks to improve upon, *SHIBA* (Webb and Ree 2012) has demonstrated that relying on a rejection step for complex biogeographic inference isn’t a trivial task and can be computationally very inefficient.

#### SSE models

State Speciation and Extinction models integrate the effects of life-history traits (e.g. morphological, geographic) along with the branching relationships on the phylogeny to infer the rate of species diversification (Maddison et al. 2007). Specifically, the *GeoSSE* model comprehensively extends the *DEC* framework by inferring the joint history of geographic ranges and species diversification (Goldberg et al. 2011). For two areas A and B, *GeoSSE* defines seven free parameters: the rates of within-area speciation in A and B, the rates of extinction in A and B, the rate of between-area vicariance, and the rates of dispersal from A to B, and from B to A (see Fig. 1 in Goldberg et al. 2011). While such fine-scale modelling of processes evidently restricts the use of *GeoSSE* to small-sized phylogenies, biogeographic processes such as *RAC* (**Fig. 2**) are absent from this framework. Nevertheless, *GeoSSE* (and other *SSE* models) can accommodate for variable diversification rates along the phylogeny even though geographic time stratification or variability such as an EGC is still out of its reach (*i.e*. for more than two areas). A fruitful avenue would be to combine the advantages of the *DEC* and *SSE* models together for understanding biogeographic patterns. In a recent study, Rolland et al. (2015) apply both approaches for studying the latitudinal diversity gradient of the Carnivora. Their results highlight the complementary nature of the *DEC* and *SSE* approaches by obtaining concordant results for the dispersal rates from temperate to tropical biomes along with the origin of the Carnivora.

#### DEC+J

The recently introduced *DEC+J* approach (Matzke 2014) models *jump* dispersal by adding another free parameter *J*. It relies on the idea of instantaneous reproductive isolation arising from founder events and formalizes a new range inheritance scenario (*i.e*. a cladogenetic event) than those previously considered by the *DEC* model. It is also useful to note that while *DECX* models rapid anagenetic change (*RAC*, *Fig. 2*), the applicability of the latter overlaps with biological situations (e.g. island clades) modeled by rapid cladogenetic change as represented by the *J* parameter in the *DEC+J* model (*cf. Inferring geographic range evolution using DECX and DEC+J*).

#### Recommendations

Since the conceptual gap between models of character and geographic evolution was bridged (Ree et al. 2005; Ree and Smith, 2008), there has been a notable increase in the number of probabilistic approaches inferring ancestral ranges using phylogenies. As a result, it has been challenging to keep track of all the methodological developments and equally be able to select an approach based on its merits. This has also typically resulted in peer-review pressure that strongly favours the use of the latest methodological development. We would like to take this opportunity to emphasize our opinion, that the choice of a set of methods (e.g. for model testing) should primarily depend upon the biological question along with the type and quality of data at hand. As a second objective, it is preferable to start with parsimonious models, in terms of free parameters, before any further complexification.

### Inferring geographic range evolution using DEC and DECX

In this paper, taking the cue from the *DEC* approach, we have introduced the *DECX* model by accommodating for additional anagenetic and cladogenetic events. Subsequently, one of the most surprising findings of our study is that *DEC* is a relatively robust model of range inference and able to infer most of the ancestral states even in cases when the true model is very different (**Table 2**). Another revealing fact is that the assumption of vicariance events at cladogenesis does not considerably alter the inferences from those of the *DEC* model. While there are some differences in the results obtained with the *DEC* and *DEC-VI* models, we might have identified one potential shortcoming of our simulation study when comparing these two models. From **Fig. 1**, it is obvious that a vicariance event (*VI*), different from alloperipatric speciation, is only possible when an ancestral area consists of at least four areas. While the total number of areas in our simulation study was eight, but given how our EGC was defined (**Fig. 2**), ancestral ranges were constrained to have no more than five areas over all the 480,000 inferences. It would thus be desirable to test specifically for the effects of vicariant events when allowing for bigger ancestral ranges in a future simulation study. However, note that the total number of ranges considered in our simulation study is larger than all other previous studies (except Landis et al. 2013). Finally, while the *RAC* component in the *DECX* model has considerably lower likelihoods when it is the true model, it only fares slightly better when *DEC* is the true model. Despite lower likelihoods, the *DECX* model introduces a lot more bias in the estimates of dispersal.

### Inferring geographic range evolution using DECX and DEC+J

It is important to understand that *DEC+J* (Matzke 2014) does not model founder-event speciation *per se*, in the sense of creating new lineages, for speciation events and their timing is information already available from the phylogeny. The *j* parameter is also not dependent on the size of the ancestral range before splitting into daughter ranges and neither does the distance from a given parent to any eventual daughter range matter. It is thus incorrect to state that *DEC+J* models rare or long-distance dispersal events. However, it is correct to state that *DEC+J* models the *relative preference* of founder events associated with speciation events already defined by the topology. While founder-event speciation in itself is not an issue of contention among biogeographers, in the following we contend that the relative nature of the *J* process conflates the role of the other two companion biogeographic processes that are dispersal and local extinction.

Inference under the *DEC+J* (Matzke 2014) often resembles that of *range shifts* or *jumps* as modelled by the *RAC* component by the *DECX*model in this paper. Our premise for modeling *RAC* (**Fig. 2**) as an anagenetic process was to account for the ensuing evolutionary cost required for any such change as a function of time. Thus, inferring range evolution using *RAC* necessarily updates the two degrees of freedom of the *DECX* model represented by the rates of dispersal and local extinction between and within areas respectively. *RAC* is thus a time-based process, fully integrated into the underlying CTMC and plays a role in the evolution of ranges along all branch lengths of the phylogeny. The jump process, represented by the *J* parameter, depends uniquely on the branching events, models instantaneous reproductive isolation and tends to explain all of the information contained in the spatial ranges of the extant species *at the expense of* the processes of dispersal and local extinction.

On an empirical note (*cf. Results: Empirical analysis of island clades*), *DEC+J* dominates in terms of model performance among the weakly constrained analyses (15/19), though a closer look reveals a more intriguing feature of the model. For all of the 19 weakly constrained analyses, *DEC+J* consistently infers zero levels of extinction, thus rendering a degree of freedom of the model redundant (Online **Appendix 8** available at https://goo.gl/oXDdJz). Except for 4 weakly constrained analyses (“*Honeycreepers_M0_unconstrained”, “Silversword_M0_unconstrained*”, “*Microlophus_M1_2areas*”, and *“Vireya_M1_2areas*”), *DEC+J* consistently infers zero levels of dispersal. Thus, among the 15 weakly constrained analyses where *DEC+J* fares better, 12 of them are inferred on the basis of a single degree of freedom. In these cases all range evolution is channelled uniquely through the cladogenetic events of sympatry, peripatry and founder jumps. **Table 3** and **Appendix 5** (available online at https://goo.gl/oXDdJz) further indicate that *DEC+J* infers non-zero dispersal (preferentially) and extinction in the presence of time-stratified constraints. For weakly constrained scenarios, the inference of dispersal and extinction is facilitated by the presence of non-endemic extant tips on the phylogeny.

In general, time-stratified constraints provide for a more complex comparison between *DECX* and *DEC+J* as the latter can be subdivided into the presence or absence of an EGC. While we do not delve any further into this detail, it suffices to know that an EGC greatly enhances the performance of *DECX* with respect to the *DEC+J* model. Finally, a fair comparison between the *DECX* and *DEC+J* models would need to include a PODs study, and the incorporation of the model constraints discussed above, which we consider to be out of scope for this paper.

### Difficulties with the empirical analysis of island clades (Matzke 2014)

The central tenet of the *DEC+J* model lies in its successful application to a notable compilation of island clades. Here, we briefly bring to the notice of the interested reader some of the difficulties we encountered while analysing the entire dataset consisting of 13 island clades from Matzke (2014, *SuppData2_all_BioGeoBEARS_inferences* available online at http://datadrvad.Org/resource/doi:10.5061/drvad.2mclt). **Table 3** mentions 9 analyses of Matzke (2014, Table 2 and Supplementary Table 1) which were either missing (*Nesophrosyne* and *Cyrtandra*) or which were conflicting (*Megalagrion5*) with the results from the aforementioned supplementary data. Also, a detailed comparison of inferred dispersal and extinction rates between the *DECX* and *DEC+J* models was hampered by an incorrect reporting of results in Matzke (2014, Supplementary Figure 9). We thus provide the inferred dispersal and extinction rates for the analyses defined in Matzke (2014, Table 2) in an accessible format (Online **Appendix 8** available at https://goo.gl/oXDdJz). Finally, we wish to point out a disagreement in the likelihoods obtained under the *BioGeoBEARS-DEC* and C++ *DEC* models, especially under time-stratified constraints and which will require further investigation of the respective implementations.

Given the inconsistencies that we have chanced upon and in the interest of scientific rigour, we warmly invite the concerned author (Matzke 2014) to issue a comprehensive *Corrigenda*, however belated, and preferably in the very journal it has been published. We also take this opportunity to commend the sole author for the compilation of this impressive volume of necessary and arduous research and for promoting the need of a rigorous model choice in the field of historical biogeography.

### A review of the DECX framework

Developed over an existing C++ framework (Smith 2009), *DECX* shares all the attributes of *DEC* in terms of model flexibility and is computationally very efficient. For instance, analyzing the amphibian tree containing 3309 species for the **M0**, **M1** and **M2** models (**Table 4**) took respectively 184, 166 and 508 seconds. This efficiency helps immensely in accounting for phylogenetic and divergence time uncertainty, as has been previously demonstrated (Smith 2009), for upcoming large phylogenies where this problem is exacerbated.

#### Evolving geological connectivity

Starting with Ree and Smith (2008), the incorporation of varying dispersal opportunity over time has been greatly facilitated by adopting a CTMC framework and by allowing the user-specified dispersal matrix **D** to parametrize the Markov transition probabilities. Consequently, a drawback appears when constructing constructing complex biogeographic scenarios and relying only on the **D** matrix as has already been highlighted by Buerki et al. (2011, p. 546). By viewing connectivity between areas as an evolving graph (**Fig. 3.**) over time intervals we have at once overcome a methodological and a conceptual issue, the latter helping us better specify the *spatial context* and thus account for an additional cladogenetic event, namely vicariance (**Fig. 2.**). The other advantage of implementing an EGC is its increased computational efficiency as the size of the **Q** matrix for every time interval is now conditioned by the respective adjacency matrix.

#### Fossil tips

Branch lengths can have a significant impact on the reconstruction of ancestral states (Cusimano and Renner 2014). Thus, the incorporation of fossil lineages along with contemporaneous lineages in a biogeographic study can also have a profound effect on the conclusions, particularly when the fossil ranges do not overlap with the extant ranges (Crisp et al. 2011). Indeed, previous studies have used the *DEC* approach to study the historical biogeography of extant groups by including extinct lineages such as the cypress trees (Mao et al. 2012) or the palpimanoid spiders (Wood et al. 2013). Some have also attempted to estimate historical biogeography of completely extinct groups such as tyrannosaurs (Loewen et al. 2013). While these studies have pioneered the use of fossil tips for biogeographic analyses, a main issue has been overlooked when using the *DEC* approach, which is the absence of an explicit method for accounting non-ultrametric trees (*cf. Methods: Empirical analysis of palpimanoids*).

When correctly accounting for the branch lengths of extinct lineages we would intuitively expect ancestral nodes neighbouring the fossil branching points to be influential in two ways as a result of the Felsenstein’s pruning algorithm. First, conditional likelihoods at nodes immediately ancestral to the fossil branching points would register a stronger probability for those states of the CTMC (or species ranges) overlapping with the fossil states. Also, the shorter the fossil branch lengths (or the earlier the fossil went extinct), the stronger would be the propagation of this signal. Second, conditional likelihoods for nodes immediately descendant of the fossil branching points would register a relative decrease in the biogeographic signal being propagated from the extant tips. As expected, our C++ *DEC* implementation did not infer the same ancestral areas as a *classic DEC* model for the palpimanoid spiders because the latter forcefully renders the phylogenies to be ultrametric thereby downing the biogeographic signal from the fossil tips (**Fig. 7.**). Though parameter inferences are typically biased under DEC based models, they can still be used to evaluate the relative importance of the underlying processes between two different analyses. For instance Wood et al. (2013) found a smaller extinction rate with respect to dispersal while we, using our C++ *DEC*, obtain an extinction rate on par with dispersal (*cf. Results: Empirical analysis of palpimanoids*). A further analysis using *DECX*, reflects the the importance of the *RAC* process as we obtain a higher extinction rate than dispersal.

#### Open questions

The improvements presented in this paper do leave questions unanswered and avenues unexplored and we take note of some of them in the following. Here, *RAC* is implemented as a single component, which under a clear model choice framework (Matzke 2014) would warrant for a finer modelling of its components. It has been recognized that the dispersal matrices **D** used in *DEC* to specify biotic movement between areas are subjective and error prone. Several papers have thus proposed biogeographic models based on distance where the rate of dispersal between any two areas is a decreasing function of the distance separating these areas (Webb and Ree 2012; Landis et al. 2013). Other possibilities include the scaling of the dispersal rate matrix based on physiological impact induced by the warming and cooling of the climate for various time intervals (Condamine et al. 2013). Also, the rate of biogeographic evolution, especially for large phylogenies, can be relaxed and allowed to be varied for different parts of the tree. For instance, one can argue that within Amphibian phylogeny the Gymnophiona (restricted to equatorial tropics) may not have the same dispersal ability as the other clades (e.g. Anura or Caudata). Finally, trees with fossil tips need to be further explored using a PODs study which would help us understand quantitatively asses the influence of fossil information on biogeographic inferences.

### Conclusions

It is not an understatement to say that we are currently in an era of integrative biogeography. While, analytical challenges remain with the increase in large phylogenies, comprehensive geographic sampling of clades and the integration of extrinsic information, the integrative nature of current progress is inspiring. As an example of the latter, a framework for the biogeographic calibration of the molecular clock has been proposed (Ho et al. 2015). Following which, Landis (2015) has proposed the first method for biogeographic dating. In a similar vein, integrating multiple line of evidence is becoming increasingly possible for recent (Metcalf et al. 2014) to deeper timescales (Meseguer et al. 2015). Thus, we cannot but agree that, “*The future of biogeography is bright and filled with exciting challenges and opportunities*” (Wen et al. 2013).

As exemplified by *DIVA*, *DEC* and variants, parametric biogeography has certainly hastened the arrival of more integrative approaches. For instance, it is now possible to test biogeographic scenarios and tease apart the role and dominance of vicariance over classic dispersal via range expansion or the relative importance of jump events during cladogenesis, as well as the role of local extinction (as inferred from fossils) in shaping biodiversity over space and time. The basic statistical flexibility of *DEC* model (Ree and Smith, 2008) has made it a relevant approach for hypothesis testing and model comparison. This paper proves once again that it still has the potential to improve upon biogeographic inferences.

## Acknowledgements

This work is based on a long-lasting collaboration initiated during the first post-doc fellowship C.R.B. and the PhD thesis of F.L.C. We warmly thank all people who discussed our ideas and projects in the different labs (e.g. Andrea Meseguer, Frédéric Delsuc, Gael Kergoat, Raphael Leblois, Mike Hickerson). They kindly pushed us to publish this work, and contribute to our ideas. A special thanks goes to Julien Veyssier for making DECX accessible to all interested users by helping us create binaries for various platforms.

